# Leucine Regulates Neuronal Health During Nutrient Deprivation in *Caenorhabditis elegans*

**DOI:** 10.64898/2026.09.04.748567

**Authors:** Muraleedharan Sudhanand, Akankshya Sahu, Janhavi Desai, Urmi Bandyopadhyay, Abhishek Bhattacharya

## Abstract

Alterations in nutrient availability induce changes in neuronal structure and connectivity across species, ultimately leading to nervous system plasticity and behavioural changes. However, it remains unclear how extended periods of acute nutrient deprivation affect nervous system integrity and function. Using the nematode *Caenorhabditis elegans*, we uncovered that prolonged macronutrient deprivation profoundly impacts neuronal health. Across developmental stages, multiple sensory neuron classes in the *C. elegans* nervous system, including polymodal IL2, volatile odor-sensing AWB and AWC, and CO_2_-sensing BAG neurons, display extensive dendritic and axonal blebbing, severely disorganized dendritic, axonal, and cell body morphologies, and severely disrupted chemical and electrical synaptic organization during prolonged nutrient deprivation. The severity of these defects increases progressively with the duration of nutrient deprivation, ultimately affecting animal behaviours. Our results show that the availability of a particular macronutrient, specifically the branched-chain amino acid leucine alone, in the absence of any other macronutrients, although insufficient to support animal growth, is sufficient to completely prevent nutrient-deprivation-mediated neuronal damage and the decline in animal behavioural responses. Moreover, leucine supplementation alone is sufficient to reverse nutrient-deprivation-mediated neuronal abnormalities. These findings suggest that the availability of leucine, rather than absolute caloric intake, is crucial for maintaining neuronal health and function under adverse conditions.

## INTRODUCTION

Animals across species experience variability in nutrient availability. Alterations in the nervous system and behaviours in response to changing nutrient availability are essential for survival and reproductive success in complex environments. Starvation-induced behavioral alterations have been observed in both vertebrates and invertebrates, including nematodes^1–10^. Animals across species also experience extended periods of acute starvation and exhibit distinct strategies to adapt to adverse environmental conditions and increase their chances of survival. Many species enter periods of hibernation or aestivation to slow down their metabolism to ultimately increase their chances of survival^11–13^4 (ref). Nematodes, including *C. elegans*, enter reversible developmental arrest or diapause stages under nutrient deprivation to increase their survival^13,14^. The nervous system in hibernation and diapause stages has been shown to undergo plastic changes^2,15–18^. Apart from environmental nutrient deprivation, metabolic disorders, aberrant cellular nutrient sensing, or reduced catabolic capacity in specific disease contexts, including aggregative neurodegenerative disorders, potentially generate prolonged local imbalance in macronutrient availability. However, how nervous systems respond to this prolonged nutrient imbalance remains to be fully understood.

*C. elegans,* with a well-characterized nervous system and well-characterized behavioural repertoire, provides a good model system to address these questions. Moreover, nutrient deprivation at distinct developmental stages leads to developmental or diapause arrest in *C. elegans*^13,14^. These diapause stages provide good models for understanding how prolonged nutrient deprivation regulates nervous system health and function. It has also been shown that the nervous system of *C. elegans* exhibits wide-ranging structural and functional plasticity in response to nutrient deprivation across life stages^2,16^. During starvation, *C. elegans* and other nematodes exhibit altered chemotaxis towards environmental CO_2_ across life stages, which has been proposed to be associated with food- and host-seeking in parasitic nematodes ^10,19–22^. During the starvation-induced dauer diapause stage, *C. elegans* show an altered dispersal behaviour, called nictation^12,13^. The IL2 class of polymodal ciliated sensory neurons undergo extensive dendritic arborization when animals enter the dauer diapause stage and regulate the dauer-stage-specific nictation behaviour^12,23^. Starvation-mediated changes in dendritic structures have also been shown in the fruit fly *Drosophila* and in the mammalian brain^24,25^.

We uncovered that prolonged nutrient deprivation leads to progressive neuronal abnormalities, including severely disorganized dendritic, axonal, and cell body morphologies, and an overall disruption of hardwired chemical and electrical synaptic connectivity across multiple neuron types and life stages. These neuronal defects lead to wide-ranging deficiencies in chemotactic behaviours of the animal during prolonged starvation. These neuronal defects can be reversed upon feeding. By dissecting the requirements for specific macronutrients in protecting neuronal health, we identified that the most abundant essential branched-chain amino acid, leucine, plays a crucial role in maintaining neuronal health. Supplementation with leucine alone, without other macronutrients, is sufficient to maintain neuronal morphological and connectivity features, thereby maintaining chemotactic behaviour of the animals during prolonged starvation. Moreover, leucine supplementation alone is sufficient to reverse prolonged starvation-mediated neuronal damages. Altogether, our findings identify a crucial neuroprotective role of leucine during prolonged nutrient deprivation.

## RESULTS

### Nutrient deprivation progressively affects the morphology of IL2 polymodal sensory neurons

*C. elegans* larvae, when hatched in the absence of food, undergo a well-characterized early-life developmental arrest, namely L1 arrest^26^. Arresting newly hatched C. elegans larvae at the L1 stage, followed by feeding them *E. coli OP50* to initiate synchronized reproductive development, is routinely used to obtain synchronized populations of animals^26^, which have formed the basis of numerous studies over the years. However, how this early-life nutrient deprivation affects neuronal health and function remains less well understood. To address this question, we investigated the IL2 class ciliated sensory neurons during the L1 arrest stage. Six IL2 neurons have their cell bodies arranged in a circle around the body axis, anterior to the nerve ring. Each extends an unbranched anterior dendrite that leads to the sensory cilia at the nose, and a single posterior axon that fasciculates in the nerve ring and forms synapses **(Figure 1A)**. Surprisingly, we observed extensive dendritic blebbing, more disorganized projections, and disorganized axonal morphology in IL2 neurons during the L1 arrest stage **(Figure 1B,C)**. Additionally, IL2 neurons display severely disorganized cell-body positioning and more rounded cell soma during the L1 arrest **(Figure 1B)**. Our results showed that IL2 neurons begin to exhibit dendritic blebbing within 12 hours of entering the L1 arrest stage, and the extent of dendritic blebbing increases progressively with the duration of the L1 arrest, peaking at 36 hours **(Figure 1B,C)**. Apart from the L1 arrest stage, nutrient deprivation before molting into the L3 and L4 larval stages also causes developmental arrest at specific checkpoint stages, namely, the L3- and L4-arrest stages, respectively^14^. Our results suggest that extensive starvation during the L4-arrest stages also causes equivalent abnormalities in IL2 neurons **(Figure 1D,E)**. However, neuronal abnormalities in IL2 neurons in L4- stage animals emerge after seven days of nutrient deprivation, compared with 12 hours of nutrient deprivation in L1-stage animals. These results together suggest that extended starvation adversely affects neural morphologies across developmental stages.

**Figure 1:**
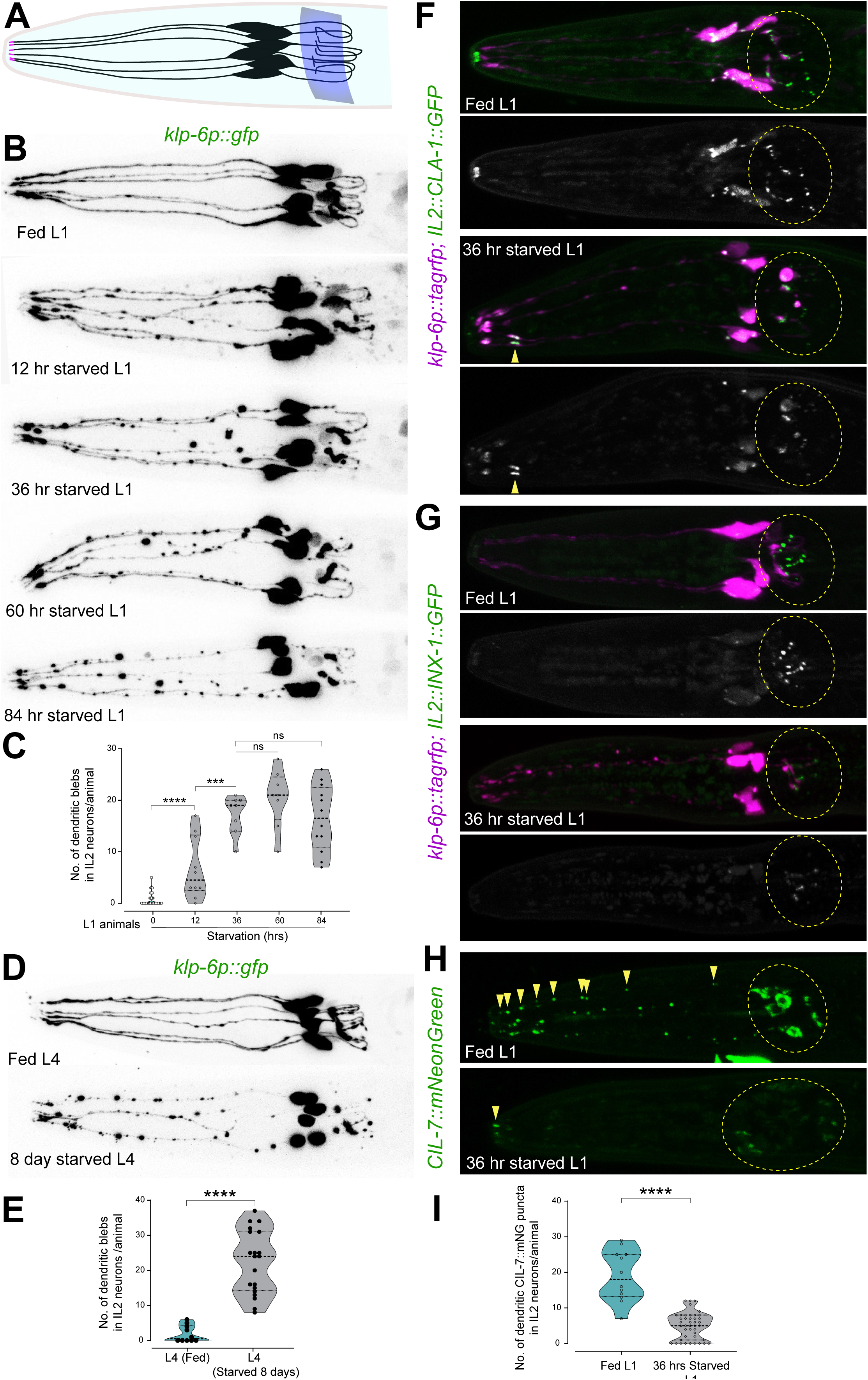
IL2 neurons in *C. elegans* show progressive defects under prolonged starvation. In all images, the anterior of the animal is on the left. (A) Schematic of IL2 neurons showing cell body positioning, anterior dendritic projections, sensory cilia (in magenta), and posterior axonal projections that lead to synapse formation in the nerve ring (blue stripe). (B) Dendrites, axons, and cell bodies of IL2 neurons in L1 stage show progressive signs of abnormalities with extended nutrient deprivation. (C) Quantification of the data shown in Panel B. Each data point represents the average number of dendritic blebs (measuring ≥0.5 µm) in IL2 neurons per animal. Violin plots represent the distribution; black horizontal lines represent the median and quartiles. Mann–Whitney U test p-values for each comparison: n.s., nonsignificant, ***p < 0.001, ****p < 0.0001. The number of dendritic blebs (measuring ≥0.5 µm) in IL2 neurons increases significantly until 36 hours of nutrient deprivation. (D) Dendrites, axons, and cell bodies of IL2 neurons at the L4 stage show significant signs of abnormalities upon 84 hours of nutrient deprivation. (E) Quantification of the data shown in Panel D. Each data point represents the average number of dendritic blebs (measuring ≥0.5 µm) in IL2 neurons per animal. Statistics as in Panel C. The number of dendritic blebs in IL2 neurons at the L4 stage increases significantly after 84 hours of nutrient deprivation. (F) IL2-specific expression of CLA-1 (CLA-1::FLPon::GFP) using cell-specific flippase-recombination showed punctate localization of CLA-1 at the IL2 active zones in the nerve ring (marked by yellow dotted circles). CLA-1 localization in IL2 active zones decreases significantly after 36 hours of nutrient deprivation. (G) IL2-specific expression of INX-1 (INX-1::FLPon::GFP) using cell-specific flippase-recombination showed punctate localization of INX-1, marking the IL2 neuron-specific electrical synapses in the nerve ring (marked by yellow dotted circles). INX-1-containing IL2 electrical synapses decrease significantly after 36 hours of nutrient deprivation. (H) Endogenously tagged CIL-7::mNG labels EVs in the IL2 neurons. Associative conditioning with IAA and heat results in significant CIL-7 localization in the IL2 cell body and EV distribution along IL2 dendrites in L1 animals. CIL-7::mNG expression in the IL2 cell body and dendrites decreases significantly after 36 hours of nutrient deprivation. (I) Quantification of the data shown in Panel H. Each data point represents the average number of CIL-7::mNG puncta in IL2 dendrites per animal. Violin plots represent the distribution; black horizontal lines represent the median and quartiles. Mann–Whitney U test p-values for the comparison: ****p < 0.0001. The number of dendritic CIL-7::mNG puncta in IL2 neurons decreases significantly after 36 hours of nutrient deprivation.

### Nutrient deprivation affects synaptic organization and extracellular vesicle distribution in IL2 neurons

IL2s are cholinergic neurons that form extensive chemical synapses in the nerve ring starting at the L1 larval stage under replete conditions, as shown by the serial-section electron-micrograph reconstruction of the nerve ring^27–29^. To understand the impact of starvation on IL2 neuronal connectivity, we examined their chemical and electrical synaptic specializations. We identified the presynaptic connectivity of IL2 neurons through cell-specific expression of endogenously green fluorescent protein (GFP)-tagged synaptic active zone marker, CLA-1/Clarinet, an ortholog of mammalian Piccolo^30^. We observed that the organization of presynaptic active zones within IL2 axons is extensively disrupted after 36 hours of L1 arrest **(Figure 1F)**.

To visualize electrical synapses in IL2 neurons, we directly observed the localization of Innexin-1 (INX-1), a component of invertebrate gap junction channels expressed in these neurons^2,31^. Using a flippase (FLP) recombinase-mediated, cell-specific GFP-tagging strategy, we found that IL2 neurons form extensive electrical synapses at the nerve ring utilizing INX-1 in L1-stage animals harbouring the *inx-1(amz08[inx-1::FLPon::gfp])* allele^32^. After 36 hours of nutrient deprivation in L1-stage animals, INX-1-mediated electrical synapses in IL2 neurons were entirely disrupted **(Figure 1G).** These results suggest that early-life nutrient deprivation significantly affects neuronal structure and synaptic connectivity. In *C. elegans* hermaphrodites, IL2 neurons are specialized to release extracellular vesicles (EVs) via sensory cilia, a process required for inter-animal communication and the transfer of long-term associative memory^33–35^. The EV release is facilitated by the conserved kinesin-3 family of motor proteins, including Kinesin-Like Protein 6 (KLP-6), which is homologous to mammalian KIF13A/B and KIF1^36,37^.

We quantified EVs in IL2 dendrites to assess EV transport in IL2 neurons during the L1 stage. It is shown that paired training of *C. elegans* with isoamyl alcohol (IAA) and heat increases EV biogenesis and release, both of which are crucial for cue-specific associative memory formation^34^. Using this associative-learning paradigm, we found that trained L1 animals that were nutrient-deprived for 36 hours exhibited a significantly lower punctate dendritic distribution of endogenously GFP-tagged CIL-7 *(cil-7(my61[cil-7::mNG])*^38^, an EV-resident myristoylated protein also necessary for EV biogenesis, compared to fed, trained L1-stage animals **(Figure 1H,I)**. Additionally, we observed reduced CIL-7::mNG accumulation in the IL2 cell bodies of starved, trained animals **(Figure 1H)**. These findings suggest that early-life nutrient deprivation impacts IL2 neuron function.

Since IL2 neuronal activity and EV release are closely linked to sensory cilia function, we tested whether early-life nutrient deprivation affects sensory cilia in these neurons. Prolonged starvation has been shown to alter primary cilium length^39^. We found that sensory cilia in IL2 neurons were significantly shorter in 36-hour-starved L1-arrested animals compared with L1 animals hatched on *E. coli* OP50. Overall, these results show that early-life nutrient deprivation significantly affects neuronal morphology, synapse organization, EV biogenesis, and distribution in IL2 neurons.

### Nutrient availability rescues neuronal abnormalities

IL2 neurons have not been observed before to exhibit dendritic blebbing and other associated morphological defects in well-fed larval and adult stages, although many of those studies began with L1 synchronization. This prompted us to test whether nutrient deprivation-induced defects in IL2 neurons during the L1 arrest stage are reversible upon feeding. To understand whether starvation-induced morphological abnormalities could be reverted upon feeding, we examined the morphology of IL2 neurons in animals that underwent 36 hours of nutrient deprivation at the L1 stage, followed by supplementation with *E. coli OP50* for varying durations **(Figure 2A)**. We found that returning to favourable environmental conditions, i.e., transfer to *E. coli OP50*, showed significant reversal of the dendritic morphology within 6 hours, and was completely reversed within 12 hours **(Figure 2A,B)**. We found that 12-hour supplementation with *E. coli OP50* rescued IL2 neurons in both solid NGM plates and in liquid growth culture conditions **(Figure S1A)**. Moreover, we found that the disrupted, discrete localization of CLA-1::GFP in pre-synaptic active zones, reduced punctate localization of electrical synaptic marker INX-1::GFP, and reduced localization of EV marker CIL-7::GFP in 36-hour-starved L1 animals were completely recovered upon feeding on *E. coli* OP50 for 12 hours **(Figure 2C-F)**. These results suggest that the starvation-induced neuronal defects are completely reversible and therefore do not reflect permanent degenerative damage.

**Figure 2:**
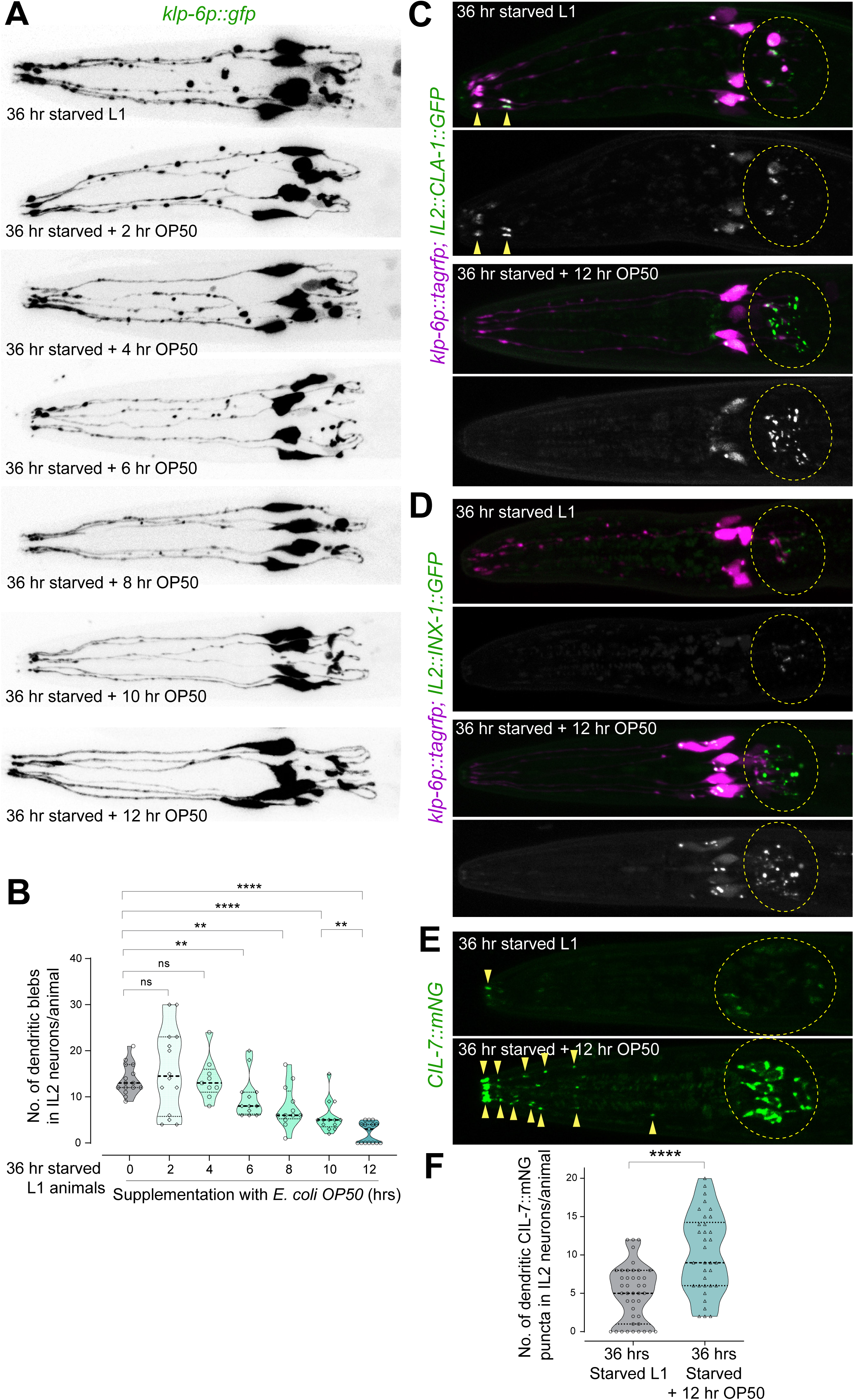
IL2 neuronal defects upon prolonged nutrient deprivation are reversible. In all images, the anterior of the animal is on the left. (A) Dendrites, axons, and cell bodies of IL2 neurons in the L1 stage show significant abnormalities after 36 hours of nutrient deprivation. These defects are progressively rescued within 12 hours of exposure to *E. coli OP50*. (B) Quantification of the data shown in Panel A. Each data point represents the average number of dendritic blebs (measuring ≥0.5 µm) in IL2 neurons per animal. Violin plots represent the distribution; black horizontal lines represent the median and quartiles. Mann–Whitney U test p-values for each comparison: n.s., nonsignificant; **p < 0.01; ****p < 0.0001. The number of dendritic blebs in 36-hour-starved IL2 neurons decreases significantly within 6 hours of exposure to *E. coli OP50* and almost completely disappears within 12 hours of OP50 exposure. (C) IL2-specific expression of CLA-1 (CLA-1::FLPon::GFP) using cell-specific flippase-recombination showed punctate localization of CLA-1 at the IL2 active zones in the nerve ring (marked by yellow dotted circles). CLA-1 localization at active zones within 36-hour-starved IL2s increases significantly within 12 hours of exposure to *E. coli OP50*. (D) IL2-specific expression of INX-1 (INX-1::FLPon::GFP) using cell-specific flippase-recombination showed punctate localization of INX-1, marking the IL2 neuron-specific electrical synapses in the nerve ring (marked by yellow dotted circles). INX-1-containing electrical synapses within 36-hour-starved IL2 neurons increase significantly within 12 hours of exposure to *E. coli OP50*. (E) CIL-7::mNG labels EVs in the IL2 neurons of L1 animals that have been trained with IAA and heat (associative conditioning). CIL-7::mNG expression in the cell body and dendrites of 36-hour-starved IL2s increases significantly within 12 hours of exposure to *E. coli OP50*. (F) Quantification of the data shown in Panel E. Each data point represents the average number of CIL-7::mNG puncta in IL2 dendrites per animal. Violin plots show the distribution; black horizontal lines indicate the median and quartiles. Mann– Whitney U test p-value for the comparison: ****p < 0.0001. The number of dendritic CIL-7::mNG puncta in 36-hour-starved IL2 neurons increases significantly after 12 hours of exposure to *E. coli OP50*.

### Availability of essential amino acids rescues neuronal abnormalities

The reversibility of IL2 neuronal defects after supplementation with *E. coli* OP50 prompted us to further investigate whether (a) the availability of macronutrients is necessary to rescue IL2 neuronal health, or alternatively (b) the improved neuronal health is related to growth associated with feeding for 12 hours. To address this, we took advantage of a previous report showing that dietary supplementation with palmitic acid alone is sufficient to initiate early postembryonic development in L1-arrested animals, even in the absence of other macronutrients^40^. Our results showed that 12-hour supplementation with 1 mM palmitic acid alone after 36 hours of L1 arrest was insufficient to rescue IL2 cell-body positioning, cell soma shape, dendritic blebbing, and axonal morphology, although it initiated growth in L1-arrested animals as observed by an increase in their body length **(Figure 3A,B)**. To test whether supplementation with other macronutrients alone is sufficient to rescue IL2 neuronal health during L1 arrest, we supplemented 36-hour L1-arrested animals with 30 mM glucose and a total amino acid cocktail (Total-AA), both of which were shown to be insufficient to initiate early postembryonic development^40^. Our results suggested that 12-hour supplementation with the Total-AA cocktail after 36 hours of L1 arrest was sufficient to rescue IL2 neuronal defects, whereas glucose supplementation showed no recovery of neuronal health **(Figure 3A-D)**. These results suggested that IL2 neuronal health during the early life stages is uncoupled from growth but linked to the availability of particular macronutrients.

**Figure 3:**
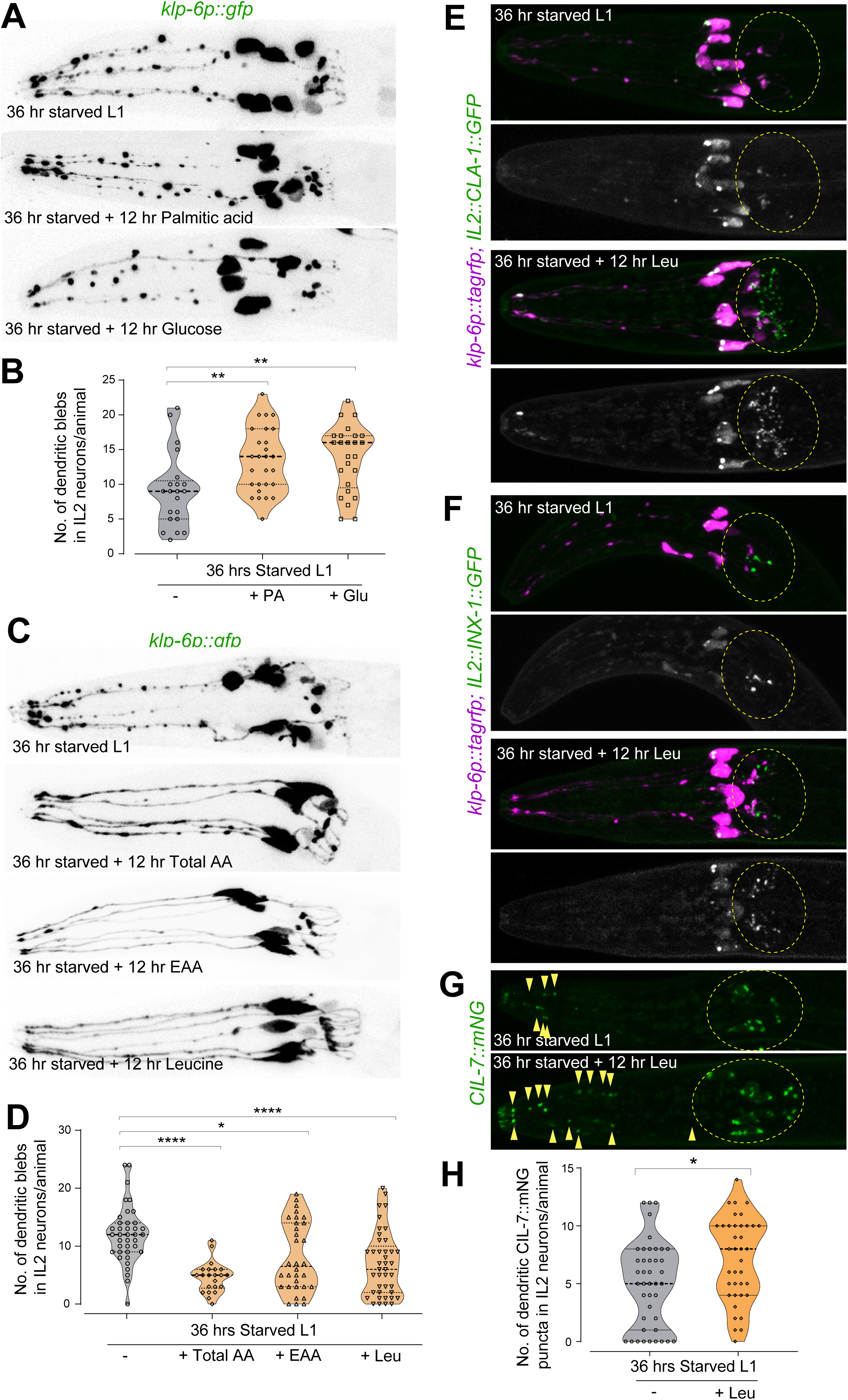
Leucine supplementation rescues prolonged nutrient deprivation-mediated IL2 neuronal defects. In all images, the anterior of the animal is on the left. (B, D, H) Each data point represents the average number of dendritic blebs (measuring ≥0.5 µm) (B and D) and dendritic CIL-7::mNG puncta (H) in IL2 neurons per animal. Violin plots represent the distribution; black horizontal lines represent the median and quartiles. Mann–Whitney U test p-values for each comparison: n.s., nonsignificant; *p < 0.05; **p < 0.01; ***p < 0.001; ****p < 0.0001. (A) Supplementation with palmitic acid and glucose fails to rescue dendritic, axonal, and cell body defects in 36-hour-starved IL2 neurons. (B) Quantification of the data shown in Panel A. The number of dendritic blebs in IL2 neurons starved for 36 hours further increased after 12 hours of supplementation with either palmitic acid or glucose. (C) Supplementation with total amino acids (AA), essential amino acid (EAA) cocktails, or leucine alone for 12 hours significantly rescued dendritic, axonal, and cell body defects in 36-hour-starved IL2 neurons. (D) Quantification of the data shown in Panel C. The number of dendritic blebs in IL2 neurons starved for 36 hours was significantly reduced after 12 hours of supplementation with total amino acids (AA), essential amino acid (EAA) cocktails, or leucine alone. (E) IL2-specific expression of CLA-1 (CLA-1::FLPon::GFP) using cell-specific flippase-recombination showed punctate localization of CLA-1 at the IL2 active zones in the nerve ring (marked by yellow dotted circles). CLA-1 localization at active zones in IL2 neurons starved for 36 hours increased significantly after 12 hours of supplementation with leucine alone. (F) IL2-specific expression of INX-1 (INX-1::FLPon::GFP) using cell-specific flippase-recombination showed punctate localization of INX-1, marking IL2 neuron-specific electrical synapses in the nerve ring (marked by yellow dotted circles). INX-1-containing electrical synapses in IL2 neurons starved for 36 hours increased significantly after 12 hours of supplementation with leucine alone. (G) CIL-7::mNG labels EVs in the IL2 neurons of L1 animals trained with IAA and heat (associative conditioning). CIL-7::mNG expression in the cell bodies and dendrites of 36-hour-starved IL2s increases significantly after 12 hours of supplementation with leucine alone. (H) Quantification of the data shown in Panel G. The number of dendritic CIL-7::mNG puncta in 36-hour-starved IL2 neurons increased significantly after 12 hours of supplementation with leucine alone.

To determine whether the complete Total-AA cocktail or specific amino acids are required for the rescue of IL2 neuronal defects, we first supplemented with only the essential amino acid (EAA) cocktail or the non-essential amino acid (Non-EAA) cocktail. We found that 12-hour supplementation with the EAA cocktail after 36 hours of L1 arrest was sufficient to rescue IL2 neuronal defects **(Figure 3C,D).** In contrast, supplementation with the Non-EAA cocktail further exacerbated IL2 neuronal defects and significantly reduced the survivability of 36-hour L1-arrested animals **(Figure 3F,G, S2A)**. These results suggested that the availability of the total EAA cocktail or specific EAAs is essential for maintaining the health of IL2 neurons during prolonged nutrient deprivation.

### Supplementation with leucine alone is sufficient to rescue or prevent neuronal abnormalities across multiple subtypes

Within the EAA cocktail, leucine, considered the most abundant AA in protein, is linked to nutrient sensing and stress responses across species. Moreover, leucine supplementation in *C. elegans* has been shown to promote axonal outgrowth and regeneration^41^. This prompted us to investigate the effect of leucine supplementation on neuronal health in L1-arrested animals. Our results showed that leucine supplementation alone for 12 hours, in the absence of any other macronutrients, following 36 hours of L1 arrest, was sufficient to rescue defects in IL2 dendritic blebbing, dendritic morphology, axonal morphology, cell-body positioning, and cell soma shape **(Figure 3C,D)**. 12-hour leucine supplementation also rescued defects in IL2 presynaptic organization, as monitored by CLA-1 localization, INX-1 localization in electrical synapses **(Figure 3E,F)**, and CIL-7-marked EV localization in IL2 dendrites **(Figure 3G,H)**. We also found that 12 hours of supplementation with other branched-chain amino acids, isoleucine and valine, could also rescue IL2 neuronal defects in 36-hour L1-arrested animals, although to a lesser extent **(Figure S2C,D)**. To determine whether the presence of leucine alone during extended nutrient deprivation, without any other macronutrients, is sufficient to preserve neuronal health, we examined the morphology of IL2 neurons in L1 animals maintained in the presence or absence of leucine. To our surprise, the presence of leucine alone was sufficient to completely maintain neuronal integrity in 84-hour-starved L1 animals **(Figure S2D)**. These results suggest that the availability of leucine and other branched-chain amino acids not only prevents deterioration in neuronal health and connectivity features of IL2 neurons during early-life nutrient deprivation, but also rescues neuronal health after starvation-induced damage.

### Nutrient deprivation affects morphology of multiple sensory neuron classes

We next tested whether the effect of long-term starvation on neuronal morphology and connectivity is specific to IL2 neurons. CO_2_-sensing BAG neuron pair in the head of the animal plays a crucial role in regulating CO_2_-chemotaxis behavior across nematode species. Similarly, AWB and AWC classes of chemosensory neurons are crucial for sensing various repulsive and attractive volatile odorants, respectively. All these neuron types have their cell bodies near the nerve ring, extend an unbranched anterior dendrite that leads to the sensory cilia at the nose, and posterior axons that synapse in the nerve ring^28^. We found that all these classes of neurons show extensive dendritic blebbing, disorganized dendritic and axonal projections, and abnormal soma morphology upon prolonged starvation **(Figure 4A-D)**. However, our data suggest that distinct neurons show varied responses to prolonged starvation. While AWB and AWC neurons showed extensive defects within 36 hours of nutrient deprivation, BAG neurons showed much less defects at the same point, and only showed extensive defects after 84 hours of nutrient deprivation **(Figure 4A-D)**. As seen with IL2 neurons, supplementation with leucine alone in the absence of other macronutrients completely prevented neuronal defects in AWB, AWC, and BAG neurons associated with prolonged nutrient deprivation **(Figure 4A-D).** These results suggest that leucine has a wide-ranging role in protecting neuronal health during nutrient deprivation.

**Figure 4:**
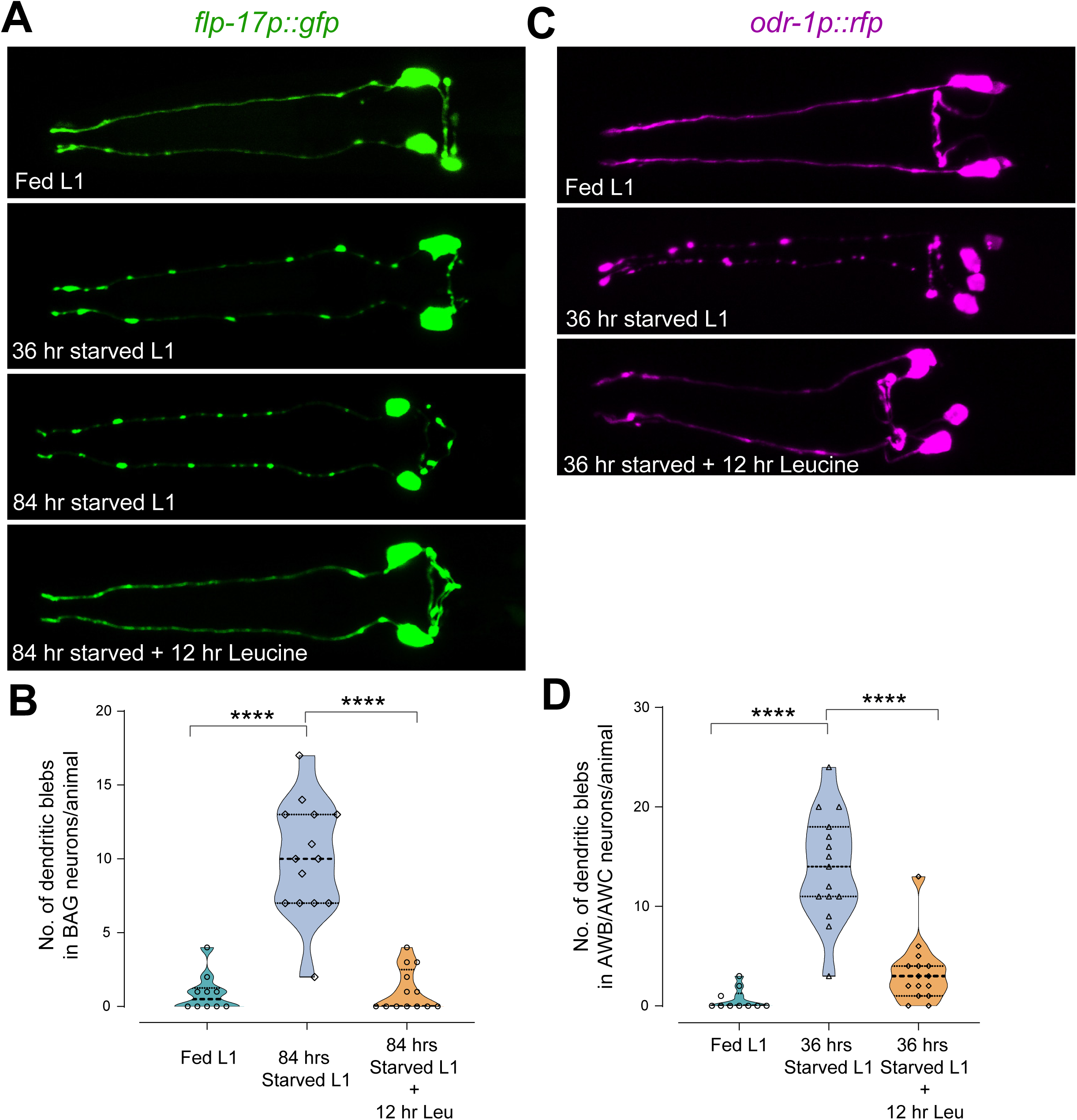
Leucine supplementation rescued prolonged nutrient-deprivation-induced defects in BAG, AWB, and AWC neurons. In all images, the anterior of the animal is on the left. (B and D) Each data point represents the average number of dendritic blebs (≥0.5 µm). Violin plots represent the distribution; black horizontal lines indicate the median and quartiles. Mann–Whitney U test p-values for each comparison: ****p < 0.0001. (A) Dendrites, axons, and cell bodies of BAG neurons (marked by flp-17p::gfp expression) after 36 and 84 hours of nutrient deprivation at the L1 stage show extensive abnormalities. Supplementation with leucine alone for 12 hours significantly rescued dendritic, axonal, and cell body defects in IL2 neurons starved for 84 hours. (B) Quantification of the data shown in Panel A. (C) Dendrites, axons, and cell bodies of AWB and AWC neurons (marked by odr-1p::rfp expression) after 36 hours of nutrient deprivation at the L1 stage show extensive abnormalities, which were significantly rescued after supplementation with leucine alone for 12 hours. (D) Quantification of the data shown in Panel C.

### Leucine is required to preserve CO_2_ chemotaxis behaviour under prolonged starvation

Wide-ranging defects in neuronal morphology and connectivity features under prolonged nutrient deprivation across neuron types prompted us to investigate the impact of nutrient deprivation on neuronal function and ultimately animal behaviour. Starvation has been shown to alter animal behaviour across species^4–9^. However, most of these studies focused on relatively short-term starvation. To understand the effect of prolonged starvation, which significantly affects neuronal health, we checked the chemotactic behaviour of *C. elegans. C. elegans* across life stages exhibit strong avoidance to environmental CO_2_ under well-fed conditions. However, animals in the dauer diapause stage and under starvation, including during the L1 arrest stage, exhibit strong attraction to CO_2_, which can be reversed to avoidance upon feeding^10,19–22^ **(Figure 5A)**. Both of these opposing valences of CO_2_ chemotaxis depend on the

**Figure 5:**
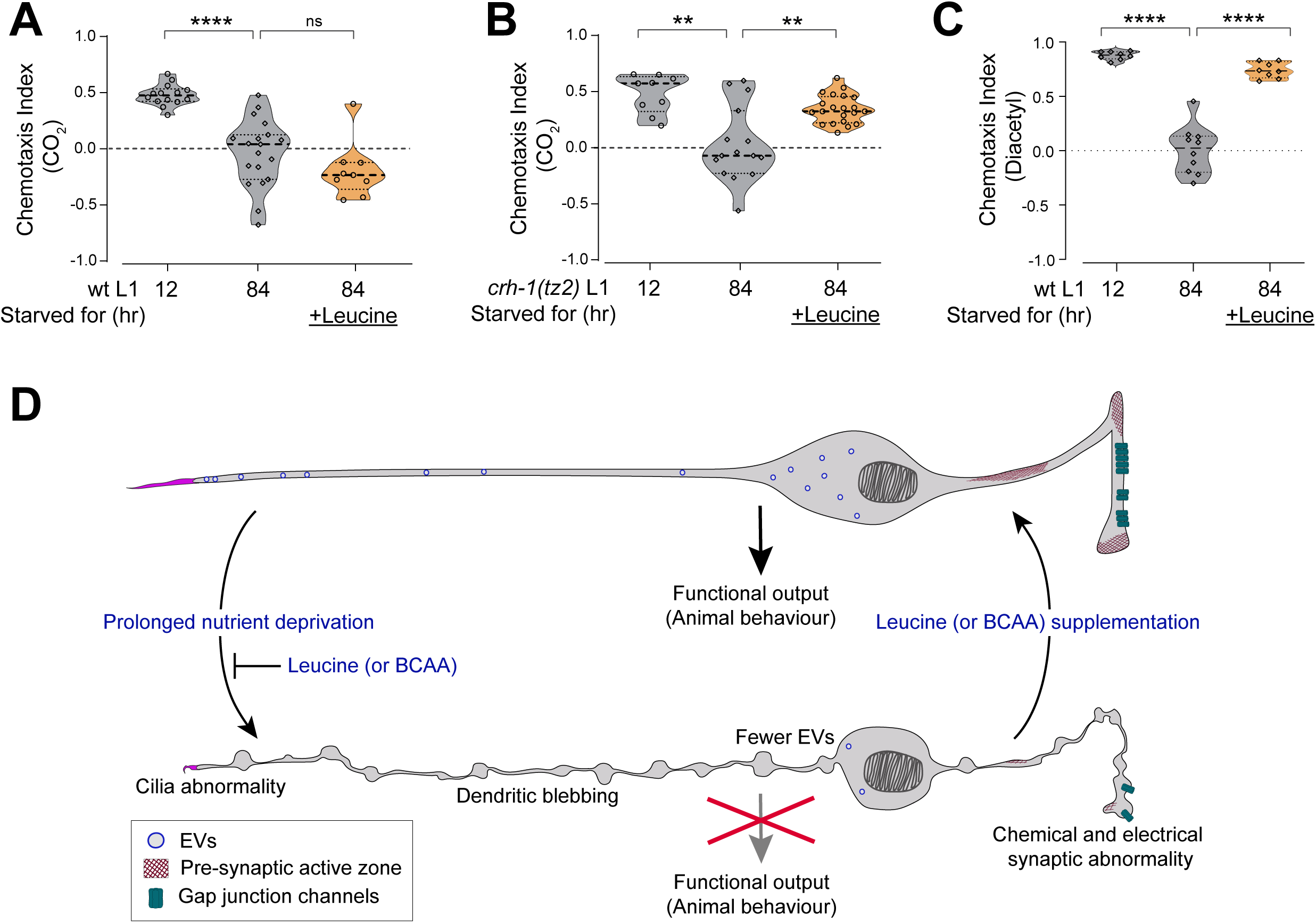
Leucine supplementation rescued prolonged nutrient-deprivation-induced defects in chemotaxis behaviours. (A-C) Each circle represents the chemotaxis index calculated from a single chemotaxis assay. Violin plots represent the distribution; black horizontal lines represent the median and quartiles. Mann–Whitney U test p-values for each comparison: n.s., nonsignificant; ****p < 0.0001. The number of animals beyond the 1 cm line from the center was counted. (A) Strong CO_2_-attraction observed in 12-hour L1-arrested animals was abolished after prolonged nutrient deprivation (84 hours), while animals supplemented with leucine during the nutrient deprivation showed mild CO_2_-avoidance. (B) *chn-1(by155)* mutant 12-hour L1-arrested animals exhibited strong CO_2_-attraction, which was abolished after prolonged nutrient deprivation (84 hours). *chn-1(by155)* mutant animals that were supplemented with leucine during the nutrient deprivation continued to show strong CO_2_-attraction. (C) Wild-type animals at the L1 stage exhibited strong attraction to diacetyl, which was abolished after prolonged nutrient deprivation (84 hours). Animals supplemented with leucine during the nutrient deprivation continued to show strong attraction to diacetyl. (D) Schematic showing that distinct morphological and connectivity features of neurons in *C. elegans* are affected by prolonged nutrient deprivation, ultimately leading to a decline in neuronal function and animal behavior. Supplementation with leucine (or other BCAAs) during nutrient deprivation prevents neuronal damage and behavioral deficits. Moreover, supplementation with leucine (or other BCAAs) is sufficient to reverse the nutrient-deprivation-induced neuronal abnormalities.

activity of BAG neurons and the BAG-expressed receptor-type guanylate cyclase, GCY-9 ^10,19^. We hypothesized that prolonged nutrient-deprivation-mediated damage to BAG neurons during the L1 stage may prevent animals from exhibiting CO₂ chemotaxis, as has been observed in animals lacking BAG neurons or that are mutant for *gcy-9,* the receptor-type guanylate cyclase expressed in BAG neurons^10,19,22^. Supporting this hypothesis, we found that nutrient deprivation for 84 hours almost completely eliminated CO₂ attraction in wild-type starved L1 animals **(Figure 5A)**. To determine whether leucine supplementation alone, in the absence of other macronutrients, could preserve chemotactic ability, we examined CO₂ chemotaxis in L1 animals deprived of all macronutrients except leucine for 84 hours. We found that leucine-supplemented L1-arrested animals showed strong CO₂ avoidance, suggesting that leucine alone could act as a feeding cue sufficient to reverse the chemotaxis valence, and leucine supplementation maintained neuronal integrity necessary for chemotaxis behaviour **(Figure 5A)**. Moreover, to further examine the neuroprotective role of leucine independent of its role as a satiety cue, which leads to reversal of CO_2_ chemotactic valence, we tested L1-stage animals mutant for *crh-1,* which encodes a homolog of the mammalian cAMP response element-binding protein (CREB). Mutation in *crh-1(tz2)* locks the CO_2_-chemotaxis valence to attraction irrespective of the feeding state^22^, allowing us to directly compare CO_2_-attraction in 84-hour nutrient-deprived L1 animals in the presence and absence of leucine supplementation. Our results suggested that leucine supplementation alone significantly improved the CO_2_-attraction in animals starved for 84 hours **(Figure 5B)**. Similarly, *C. elegans* exhibit strong attraction towards the bona fide food odor 2,3-butanedione (diacetyl), which is sensed by the AWA class of sensory neurons^42^. We found that *C. elegans* at the L1 arrest stage, which had been nutrient-deprived for 12 hours, continued to show strong attraction towards diacetyl (**Figure 5C)**. This chemotactic preference towards diacetyl was almost completely lost after 84 hours of nutrient deprivation, when most neuron types exhibited significant defects in health and connectivity **(Figure 5C)**. However, supplementation with leucine alone, in the absence of other macronutrients, rescued the decline in chemotactic preference for diacetyl **(Figure 5C)**. Together, these results suggest that leucine supplementation alone, in the absence of other macronutrients, is sufficient to maintain nervous system activity during prolonged starvation.

## DISCUSSION

Across species, animals experience extended periods of acute starvation and employ distinct strategies to endure adverse environmental conditions, thereby increasing their chances of survival. Acute starvation has also been shown to influence animal behaviours that facilitate food and mate searching, ultimately affecting the survival of the organism. In this study, we show how acute nutrient deprivation affects neuronal health and connectivity in *C. elegans* across developmental stages. Under short-term starvation, *C. elegans* alter their chemotactic and locomotory behaviours, which may facilitate their survival in natural habitats characterized by fluctuating resource availability. We show here that prolonged nutrient deprivation leads to progressive and wide-ranging abnormalities in sensory neurons of *C. elegans*. Specifically, multimodal IL2 sensory neurons, CO_2_-sensing BAG neurons, and volatile odor-sensing AWB and AWC neurons exhibit extensive neuronal damage and loss of connectivity under prolonged nutrient deprivation. However, the time scales of these damages differ among neuron types, potentially due to differences in metabolic activity and neuronal cytoskeletal organization. Our findings provide a model to understand how acute nutrient deprivation can adversely affect neuronal health.

Our work indicates that prolonged nutrient deprivation alters the clustering of presynaptic active zone proteins and gap junction channels at electrical synapses. We found that both the protein levels and the synaptic localization of CLA-1/Piccolo and the innexin, INX-1, decrease markedly during prolonged nutrient deprivation. It was shown that in the hibernating ground squirrel brain, the synaptic localization of the presynaptic marker Piccolo, the postsynaptic marker PSD95, and Synaptophysin is also severely disrupted. Future studies are required to understand how nutrient availability affects synaptic protein organization across species. Our findings also demonstrate that prolonged nutrient deprivation impairs the functional output of neural circuits. A primary function of IL2 neurons in *C. elegans* is the release EVs to mediate inter-individual communication and transfer of associative learning experiences. EV biogenesis and their dendritic transport are severely affected under prolonged starvation. Additionally, chemotaxis to various olfactory cues is significantly diminished following prolonged nutrient deprivation, underscoring the importance of maintaining neuronal health under adverse nutrient availability.

Our findings demonstrate that nutrient availability, particularly leucine and other BCAAs, induces rapid and widescale remodelling of hardwired synapses, independent of the growth of the animal. This reformation of synaptic structures occurs within hours, enabling nutrient availability to mediate alterations in neural circuits. Future studies are required to elucidate whether and how post-starvation synaptic rewiring affects neural circuit plasticity and long-term memory of starved conditions. We also found that leucine availability in the absence of other macronutrients could function as a neuroprotective agent, maintaining neuronal health, synaptic integrity, and functional output. Metabolic disorders, lysosomal dysfunction, or impaired lysosomal degradative capacity, particularly in protein-aggregative disorders, may result in a local deficiency of leucine, the most abundant amino acid. It would be interesting to investigate in the future how leucine availability affects neuronal health under these pathophysiological conditions. Supporting this hypothesis, dietary supplementation with selective EAAs has been shown to alleviate symptoms related to aging and tauopathy^43^.

Our work provides a foundation for further investigation into the mechanisms by which leucine availability mediates the maintenance or restoration of neuronal health. Leucine has been shown to upregulate the conserved mechanistic target of rapamycin complex 1 (mTORC1) pathway, thereby regulating cell growth and metabolism^44,45^. However, it remains unclear whether the mTORC1 pathway contributes to leucine-mediated regulation of neuronal health and function. It also remains to be elucidated in which tissue type leucine functions to maintain neuronal health. Finally, animals in the dauer diapause stage experience prolonged starvation, sometimes lasting up to months. Despite this, sensory neurons in *C. elegans* during the dauer diapause stage do not exhibit similar nutrient-deprivation-mediated deterioration. Future studies are required to elucidate the mechanisms by which neuronal integrity is maintained during the dauer stage.

## EXPERIMENTAL DETAILS

### *C. elegans* strains and handling

*C. elegans* strains were cultured at 22°C on nematode growth media (NGM) plates supplemented with *E. coli,* strain OP50, as a food source according to the standardized protocols^1^. *C. elegans* strain Bristol N2, was used here as the wildtype strain. For IL2 experiments, PT2519 strain, *myIs13 [klp-6p::gfp + pBS]; him-5(e1490)*^30^, was used as the control. Transgenic strains were generated using the standard microinjection technique.

A complete list of strains and transgenes used in this study is listed in **Table S1**.

### L1 synchronization and starvation assays

To obtain synchronized L1-arrested animals for all the *C. elegans* strains used in this study, gravid adults that were well-fed for at least two generations on *E. coli* OP50 were bleached according to a standardized hypochlorite bleaching protocol^46^. Eggs were hatched in M9 buffer under shaking on a tabletop shaker at 22°C. L1-arrested animals were collected at specified time intervals (12, 36, 60, and 84 hours) from the time of hatching by centrifugation at 300 x g and imaged.

### OP50 feeding for starvation-rescue assays

Synchronized L1-arrested animals maintained in M9 were collected after 36 hours of starvation, unless mentioned otherwise, by centrifugation. The collected L1s were washed 3 x times with M9. After the wash, the L1 larvae were transferred to an OP50 feeding solution and incubated on a tabletop shaker at 22 °C. The feeding solution was prepared by pelleting 5 mL of freshly grown overnight OP50 culture, then re-suspending the bacterial pellet in 10 mL of sterile M9 buffer. Animals were imaged after 12 hours of feeding, unless mentioned otherwise.

### Macronutrient supplementation

For macronutrient supplementation assays, 36-hour-starved L1-arrested animals were collected as mentioned above. The collected L1 larval population was transferred to the respective macronutrient supplementation solutions in M9, and imaged after 12 hours, unless mentioned otherwise.

Both Palmitic acid and glucose supplementation were performed as previously described in *Ruan, et al. 2024*^40^. Palmitic acid (Sigma, Cat. P5585) supplementation solution was obtained by diluting 100 mM stock solution in DMSO in M9 buffer to a final concentration of 1 mM.

The Glucose (Qualigens, Cat.15405) supplementation solution was obtained by diluting 100 mM stock solution in sterile ddH_2_0 in M9 buffer to a final concentration of 30 mM.

Amino acid stock solutions were made in sterile ddH_2_0. Amino acid supplementation solution was obtained by diluting: 100X stock solution of non-essential amino acids (Thermofisher, Cat.11140050) to 1X working concentration; 50X stock solution of Essential amino acids (Thermofisher, Cat.11130036) to 1X working concentration; leucine (Sigma Aldrich, Cat.L8912) 50 mM stock solution to 0.5 mM final concentration, methionine.

### LTAM training paradigm to study CIL-7 distribution in IL2 neurons

The Long-Term Associative Memory (LTAM) training paradigm was designed as described in *Bhar et al., 2025*^34^ with minor modifications. L1s with endogenously tagged CIL-7, *cil-7(my61[cil-7::mNG])*^38^, in all 4 conditions: fed, starved, fed after starvation, and Leu-supplemented were collected following washing and subjected to an aversive (heat; 37°C) and attractive (chemoattractant; Isoamyl alcohol) cue simultaneously. 2 minutes of exposure was followed by a rest period at 22°C for 10 minutes. This cycle was repeated 5 times, after which the worms were immediately imaged to study CIL-7 localization in IL2 neurons.

### Cloning and constructs

For the *klp-6p::TagRFP* construct, TagRFP was cloned under the *klp-6p* driver in the pPD95.75 vector backbone. *klp-6p::TagRFP* was injected at 10 ng/μl concentration to make the respective transgenes. As the IL2 cells are visible at this concentration of RFP, no other co-injection markers were used.

### CO_2_ Chemotaxis assay

CO_2_ chemotaxis assays were performed on 9cm NGM plates, as previously described^2,19,22^. A mixture of (10% CO_2_ + 20% O_2_ + 70% N_2_) was pumped through one inlet, while another mixture of (20% O_2_ + 80% N_2_) was pumped through another inlet, using programmable syringe pumps (New Era Scientific Pump Systems) to generate the CO_2_ gradient. ∼100-150 L1 animals were placed at the center of the assay plates. Assays were performed at a flow rate of 1 ml/min for 60 minutes. The number of animals that moved beyond the 1 cm line toward either the air or CO₂ side was counted to calculate the chemotaxis index (C.I.).

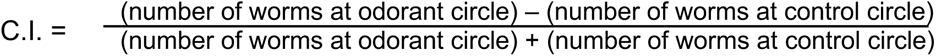

L1-arrested population was habituated on NGM agar plates for 2 hours prior to the chemotaxis assay. Results were excluded if the difference in C.I. values between pairs was ≥0.9 or if ≥5 worms failed to reach the scoring regions.

### Odortaxis assay

L1 odortaxis assays were performed on standard 9cm NGM plates. A diacetyl gradient was set by spotting 2 μL of diacetyl (1:1000 (v/v) dilution) in a spot 1 cm away from the diameter, and spotting 2 μL of ethanol (Control) at a spot exactly opposite to the diacetyl spot, and 1 cm away from the diameter. Scoring regions were the 1 cm line toward either the control half or CO₂ side. Animals in each half were counted to calculate the chemotaxis index (C.I.). ∼100-150 L1 animals were placed at the center of the assay plates and allowed to chemotax for 60 min.

The chemotaxis index (C.I.) was calculated as:

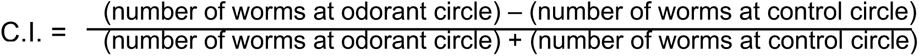

### Microscopy

Worms were anesthetized using 100 mM of sodium azide and mounted on 5% agarose pads on glass slides. Imaging was done using NIKON AX confocal laser-scanning microscopes and Olympus FV3000. Image processing and analysis were performed by scanning the full Z-stack using NIH Fiji software. Maximum-intensity projections of representative images were generated using NIH Fiji software. Figures were prepared using Adobe Photoshop 2025 and Adobe Illustrator 2025. Separate channels were usually adjusted independently using Levels and Curves in Adobe Photoshop.

### Quantification and Statistical Analysis

Although the cell body size and positioning are notable phenotypic differences between starved and supplemented/fed animals, **t**he number of blebs in the IL2 dendrites was used as the quantification strategy to compare between different groups throughout the study. To ensure consistent scoring and to exclude minor dendritic irregularities that were present across all experimental groups (starved, fed, and supplemented animals), a minimum size threshold of 0.5 µm was established for bleb identification. Only dendritic protrusions measuring ≥0.5 µm were counted as blebs for quantitative analyses.

GraphPad Prism 8.0 was used to plot graphs and perform statistical analyses. The Mann–Whitney U test (two-tailed) was used for all quantifications used in this study. Data are presented as individual bleb counts from each animal. P<0.05 was considered significant.

### Contact For Reagent and Resource Sharing

Detailed protocols are included in the article and/or supporting information. All *C. elegans* strains will be deposited at the Ceanorhabditis Genetics Center (CGC) upon publication. Request for further information will be fulfilled by the corresponding author, Abhishek Bhattacharya.

## ACKNOWLEDGMENTS

We thank Selvanayaki Eswaramoorthy and the NCBS *C. elegans* facility for assistance with worm experiments; wormbase.org and wormwiring.org for resources; Caenorhabditis Genetics Center (CGC), Arnab Mukhopadhyay, Kavita Babu, for *C. elegans* strains; members of the Bhattacharya lab for comments on this manuscript. This work was supported by DBT Wellcome Trust India Alliance (IA/I/20/2/505211) and by the Department of Atomic Energy, Government of India (Project Identification No. RTI-4018).

## AUTHOR CONTRIBUTIONS

A.B. and U.B. conceptualized the project; S.M. and A.B. designed experiments and oversaw the project; S.M., A.S., and J.D. performed experiments; S.M. and A.B. analyzed and interpreted data; A.B. acquired funding and wrote the paper.

## SUPPLEMENTARY FIGURE LEGENDS

**Figure S1: Nutrient-deprivation-induced IL2 neuronal defects are reversible after feeding in both liquid culture and NGM agar plates**

In all images, the anterior of the animal is on the left. Dendrites, axons, and cell bodies of IL2 neurons in the L1 stage show significant abnormalities after 36 hours of nutrient deprivation. These defects were rescued within 12 hours of supplementation with *E. coli OP50*, both in liquid culture and on NGM agar plates.

**Figure S2: Supplementation with BCAAs reverses the nutrient-deprivation-induced IL2 neuronal defects**

In all images, the anterior of the animal is on the left.

(B) Supplementation with Non-EAA fails to rescue dendritic, axonal, and cell body defects in 36-hour-starved IL2 neurons.

(C) Dendritic, axonal, and cell body defects in IL2 neurons starved for 36 hours were significantly rescued after 12 hours of supplementation with BCAAs, isoleucine, and valine.

(D) Quantification of the data shown in Panel B. Each data point represents the average number of dendritic blebs (≥0.5 µm) in IL2 neurons per animal. Violin plots represent the distribution; black horizontal lines represent the median and quartiles. Mann–Whitney U test p-values for each comparison: *p < 0.05; ***p < 0.001; ****p < 0.0001.

(E) In L1-stage animals starved for 84 hours, IL2 neurons exhibited extensive dendritic, axonal, and cell-body defects. Continued supplementation with leucine alone significantly reduced these defects.

