## Supplementary figures and images for "Leucine Regulates Neuronal Health During Nutrient Deprivation in *Caenorhabditis elegans*"

### Supplemental Figures

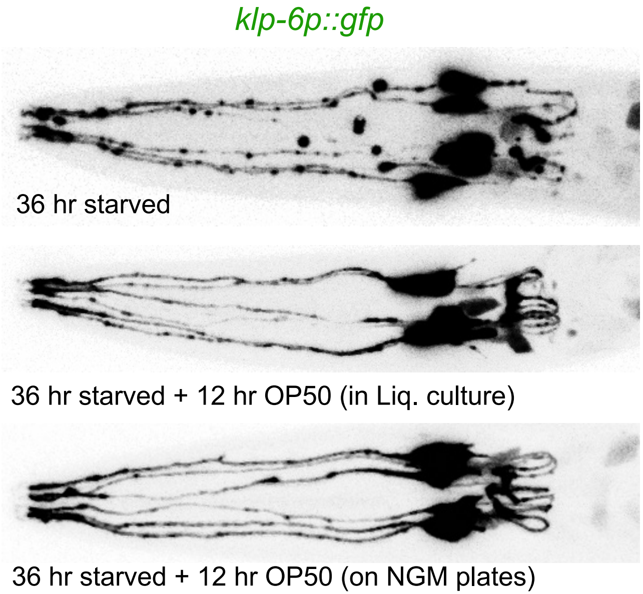

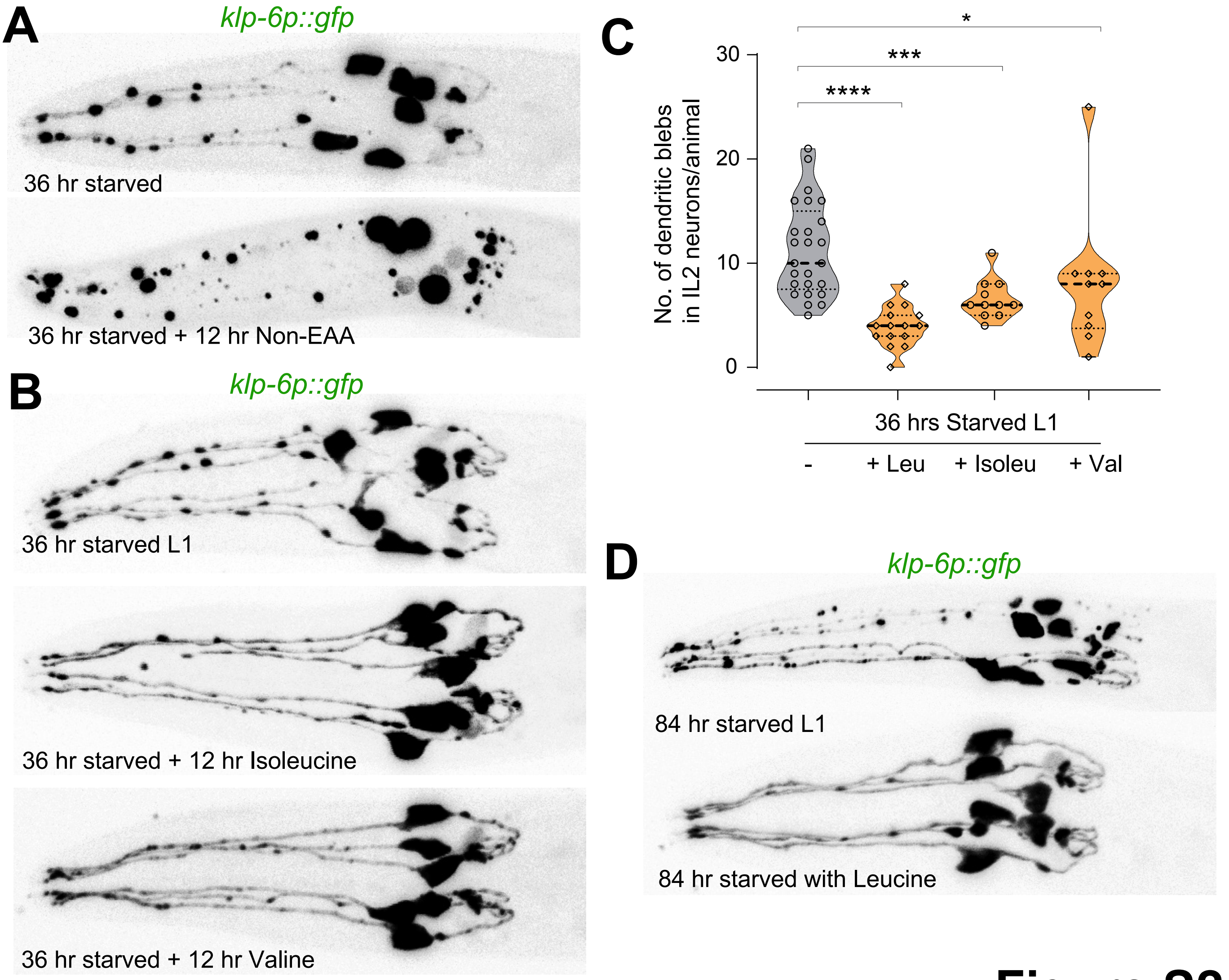
