## Supplemental Table 1 for "Leucine Regulates Neuronal Health During Nutrient Deprivation in *Caenorhabditis elegans*"

| **Strain name** | **Genotype** | **Source** |
|---|---|---|
| PT2519 | *myIs13[klp-6p::gfp + pBX]III; him-5(e1490) V* | CGC |
| PY2417 | *oyIs44[odr-1::dsRed; lin-15(+)]* | CGC |
| NY2064 | *ynIs64[flp-17p::gfp] I; him-5(e1490) V* | CGC |
| TV23058 | *unc-119(ed3)III; cla-1(wy1186[cla-1::FLPon::gfp] )IV* | Xuan et al., 2017 |
| PT3602 | *cil-7(my61[cil-7::mNG])I; him-5(e1490) V* | Wang et al, 2021 |
| ABH482 | *inx-1(amz08 [inx-1::FLPon::gfp]); amzEx147[klp-6p::Flp + klp-6p::tagRFP]* | This paper |
| ABH458 | *cla-1(wy1186[cla-1::FLPon::gfp] )IV; amzEx147[klp-6p::Flp + klp-6p::tagRFP]* | This paper |
